# Latitude delineates patterns of biogeography in terrestrial *Streptomyces*

**DOI:** 10.1101/032169

**Authors:** Mallory J Choudoir, James R Doroghazi, Daniel H Buckley

**Author notes:** Corresponding author: Daniel H Buckley School of Integrative Plant Science Bradfield Hall 705, Cornell University, Ithaca, NY 14853.

## Abstract

We examined the biogeography of *Streptomyces* at regional spatial scales to identify factors that govern patterns of microbial diversity. *Streptomyces* are spore forming filamentous bacteria which are widespread in soil. *Streptomyces* strains were isolated from perennial grass habitats sampled across a spatial scale of more than 6,000 km. Previous analysis of this geographically explicit culture collection provided evidence for a latitudinal diversity gradient in *Streptomyces* species. Here we evaluate the hypothesis that this latitudinal diversity gradient is a result of evolutionary dynamics associated with historical demographic processes. Historical demographic phenomena have genetic consequences that can be evaluated through analysis of population genetics. We applied population genetic approaches to analyze population structure in six of the most numerically abundant and geographically widespread *Streptomyces* phylogroups from our culture collection. *Streptomyces* population structure varied at regional spatial scales and allelic diversity correlated with geographic distance. In addition, allelic diversity and gene flow are partitioned by latitude. Finally, we found that nucleotide diversity within phylogroups is negatively correlated with latitude. These results indicate that phylogroup diversification is constrained by dispersal limitation at regional spatial scales and they are consistent with the hypothesis that historical demographic processes have influenced the contemporary biogeography of *Streptomyces*.

**Originality-Significance Statement:** We provide the first population genetic evidence that patterns of *Streptomyces* biogeography, which manifest in geographically explicit patterns of gene flow and a latitudinal gradient of nucleotide diversity, result from dispersal limitation and regional diversification due to drift. This contribution elucidates evolutionary processes that underlie patterns of microbial biogeography.

## Introduction

Patterns of microbial biogeography have been widely documented (Whitaker *et al*., 2003; Vos and Velicer, 2008; Bissett *et al*., 2010; Martiny *et al*., 2011; Gilbert *et al*., 2012; Hatosy *et al*., 2013), and yet we are only beginning to understand the evolutionary forces that generate and maintain these patterns. Explorations of biogeography are valuable because biogeographical patterns illustrate fundamental principles of evolution and ecology. Biogeographical patterns are ultimately governed by rates of dispersal and diversification (Martiny *et al*., 2006; Hanson *et al*., 2012). Since microbial dispersal cannot be observed directly, rates of dispersal are typically inferred from extant patterns of genetic diversity. It has been hypothesized that microbes disperse ubiquitously due to their small cell size and massive population numbers (Finaly, 2002; Finlay and Fenchel, 2004). Yet endemism and dispersal limitation have been observed for a range of microbes (Cho and Tiedge, 2000; Green and Bohannan, 2006; Telford *et al*., 2006; Boucher *et al*., 2011), and microbial dispersal limitation has been verified experimentally (Bell, 2010). Contradictory findings in the literature can be explained by at least two factors: first, dispersal constraints are likely to vary with respect to different species and habitats; and second, methods used to define units of diversity vary dramatically in their taxonomic and phylogenetic resolution. Each of these factors has been discussed previously (Hanson *et al*., 2012; Choudoir *et al*., 2012), and we will only consider them briefly here.

Patterns of microbial dispersal and gene flow appear to differ between habitats and species. At one end of the spectrum, globally widespread microbes such as *Prochlorococcus* and *Pelagibacter* show little variation in gene content between the Atlantic and Pacific Oceans suggesting that dispersal can homogenize genetic diversity in pelagic systems (Coleman and Chisholm, 2010). At the other end of the spectrum are extremophiles such as *Sulfolobus* and thermophilic *Synechococcus*, which live in island-like volcanic habitats and exhibit strong patterns of allopatric divergence resulting from dispersal limitation (Papke *et al*., 2003; Whitaker *et al*., 2003; Cadillo-Quiroz *et al*., 2012). Terrestrial microbes fall somewhere between these extremes. For example, soil dwelling microbes such as *Burkholderia pseudomallei*, *Burkholderia mallei*, and *Bacillus anthracis* exhibit biogeographical patterns governed by dispersal limitation at regional spatial scales (Kenefic *et al*., 2009; Pearson *et al*., 2009).

The phylogenetic resolution at which microbial diversity is defined can have a profound impact on our ability to discern patterns of microbial biogeography (as reviewed Hanson *et al*., 2012). Surveys of SSU rRNA genes in terrestrial habitats indicate that environmental variables including temperature (Fierer *et al*., 2009; Miller *et al*., 2009), pH (Fierer and Jackson, 2006; Lauber *et al*., 2009; Rousk *et al*., 2010), and salinity (Lozupone and Knight, 2007) are more important than geographic distance or latitude in determining spatial patterns of microbial diversity. However, SSU rRNA gene sequences have an extremely low rate of nucleotide substitution (Ochman *et al*., 1999), and microbes with similar or even identical SSU rRNA genes can have extensive genomic and ecological diversity (Welch *et al*., 2002; Jaspers and Overmann; 2004). Thus, this marker has low sensitivity for detecting neutral processes that drive patterns of biogeography, such as dispersal limitation and genetic drift (Green and Bohannan, 2006; Hanson *et al*.,. 2012). However, these neutral processes are readily explored using geographically and ecologically explicit culture collections characterized at high genetic resolution.

*Streptomyces* are ubiquitous across soil habitats, and many species are easily cultured, making this genus an excellent candidate for making a taxon-specific survey of biogeography. Furthermore, *Streptomyces* species have high rates of gene exchange both within and between species (Doroghazi and Buckley, 2010). Hence, this genus is an ideal model to explore dispersal limitations and gene flow in terrestrial systems. *Streptomyces* are gram-positive *Actinobacteria* (Kâmpfer, 2006) known for their complex developmental cycle, which entails filamentous growth and the formation of desiccation resistant spores which are readily dispersed (Keiser *et al.*, 2000). *Streptomyces* play a significant role in the terrestrial carbon cycle (McCarthy and Williams, 1992; Takasuka *et al*., 2013), represent important agricultural pathogens (Loria *et al*., 2006; Labeda, 2011), and are prolific producers of antibiotics (Watve *et al*., 2001). Despite their importance, we lack an evolutionary framework to understand *Streptomyces* biogeography.

*Streptomyces* diversity varies spatially, though the influence of geographic distance and ecological variation remains poorly resolved. Ecological adaptation constrains the environmental distribution of *Streptomyces* and their genetic and phenotypic diversity can vary in relation to soil characteristics at small spatial scale (1 m – 60 m) in prairie soils (Davelos *et al*., 2004a and 2004b), and in dune habitats (Antony-Babu *et al*., 2008). There is also evidence for dispersal limitation at very large (continental) spatial scales with endemic species observed in North America and Central Asia (Wawrik *et al*., 2007). Remarkably, genetic analysis of *Streptomyces pratensis* has revealed that strains of this species are in linkage equilibrium (i.e. random association of alleles at different loci) across a range that spans 1,000 km (Doroghazi and Buckley, 2010 and 2014). One interpretation of this finding is that *Streptomyces* are subject to widespread dispersal and unlimited gene flow at regional spatial scales. However, linkage equilibrium can also be observed for dispersal limited species that have undergone a recent historical demographic range expansion (Doroghazi and Buckley, 2010 and 2014). Demographic range expansion has previously been implicated as a factor that can explain ancestral patterns of gene exchange in *Streptomyces* clades (Andam *et al*., 2016a). Combined, these data suggest a role for both adaptive and neutral processes in governing the diversity and biogeography of *Streptomyces*, but the evolutionary interpretation of these data depends on the degree to which dispersal limitation drives patterns of diversity in *Streptomyces*.

We evaluated *Streptomyces* biogeography to explore the ecological and evolutionary forces that govern diversification within this genus. The most powerful approach for detecting evolutionary patterns that result from dispersal limitation is to examine taxon-specific biogeography patterns across ecologically similar sites. This approach controls for the effects of selection and provides a powerful test of neutral evolutionary processes (Hanson *et al*., 2012). We constructed a taxon-specific isolate collection of *Streptomyces* found in ecologically similar grassland sites spanning the United States of America. In a study of *Streptomyces* species-level diversity, which evaluated polymorphism at a single locus (*rpoB*), we observed evidence for dispersal limitation manifesting in a latitudinal diversity gradient (Andam *et al*., 2016b). The latitudinal diversity gradient is one of the earliest and most well documented patterns of biogeography (Wallace, 1878; Hillebrand, 2004), but it is essentially undocumented in terrestrial bacterial systems. We hypothesize that historical demographic phenomena associated with Pleistocene glaciation generated the latitudinal gradient of *Streptomyces* species diversity (Andam *et al*., 2016b). This hypothesis predicts that dispersal and gene flow will be limited and discontinuous across latitudes which correspond to the maximal extent of historical glaciation. The hypothesis also predicts that intra-species diversity will vary with latitude as a consequence of limited time for diversification in northern latitudes. These hypotheses are best addressed using population genetic approaches. Here we test these specific predictions by using multilocus sequence analysis (MLSA) to characterize population-level patterns of gene flow and genetic diversity within six phylogroups of *Streptomyces* which are distributed widely across the United States.

## Results

### Characterization of Streptomyces phylogroups

We identified a total of 308 isolates representing the six targeted phylogroups, and these isolates spanned 13 sites (Table 1). Strains within phylogroups share 99.4-100% SSU rRNA gene sequence identity, and all strains share greater than 97% nucleotide identity in their SSU rRNA gene sequences (Figure S1). Strains within phylogroups also share 97.6%-99.5% average nucleotide identity (ANI) across concatenated MLSA loci, and all strains share greater than 88% ANI (Figure 1, Table 2). The pattern of genetic ancestry as determined by population structure analysis (see Structure analysis, Experimental Procedures) is congruent with phylogroup boundaries (Figure S2), and this indicates that these phylogroups approximate biological populations. We observed 122 unique MLSA haplotypes, with each phylogroup represented by 13-26 haplotypes (Figure 1, Table 2). Good’s coverage for each of the six phylogroups ranges from 0.88-1.0 for individual loci and 0.94-1.0 for concatenated MLSA loci clustered at 99% nucleotide identity (Table S1). Hence, allelic diversity and nucleotide diversity are well sampled (Table S1), though unique haplotypes remain under sampled (Figure S3). The per site nucleotide diversity (π) of each phylogroup ranges from 0.0026 to 0.011 (Table 2).

**Figure 1.**
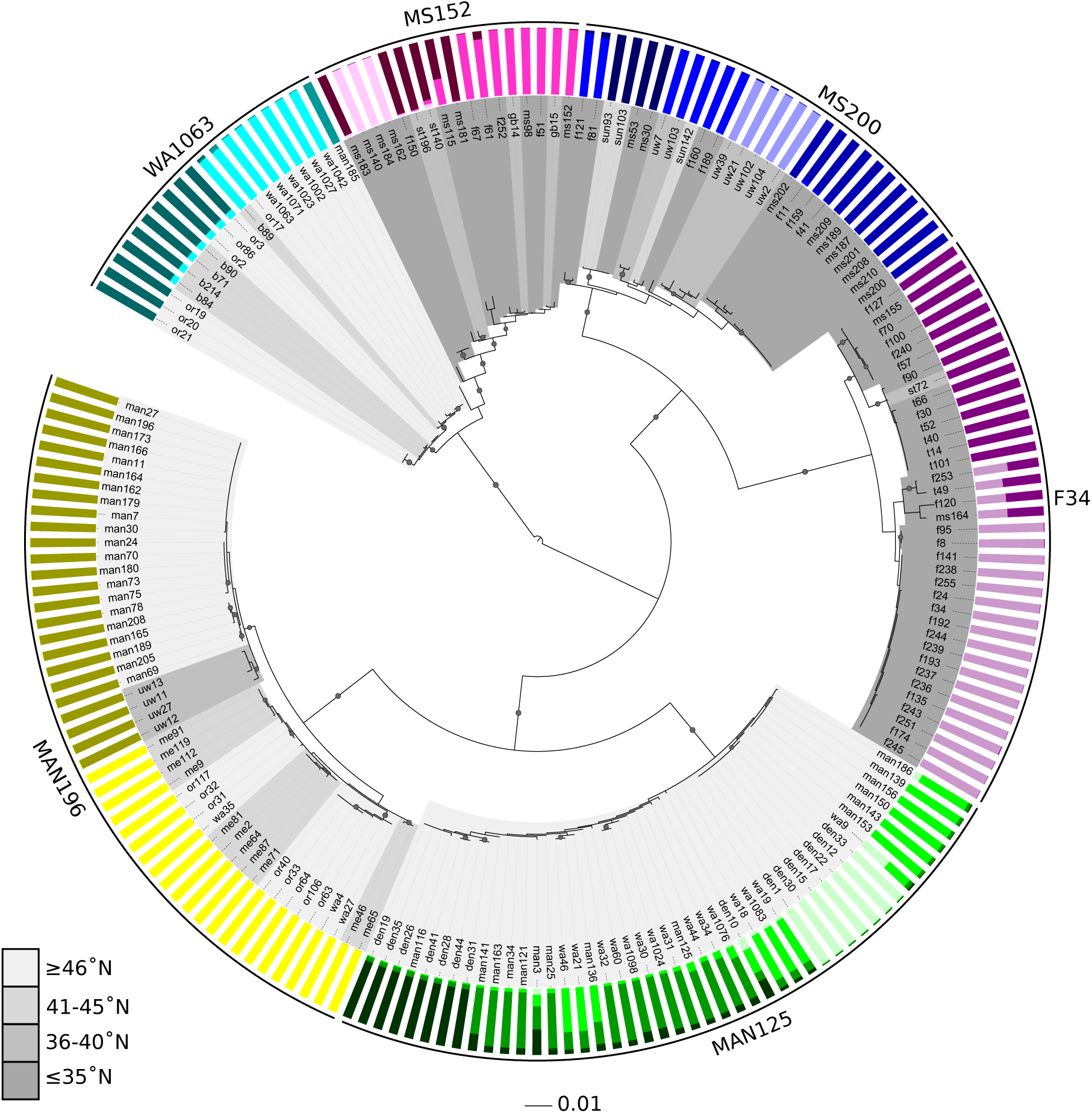
Tree was constructed from concatenated MLSA loci nucleotide sequences using maximum likelihood with a GTRGAMMA evolution model. Scale bar represents nucleotide substitutions per site. The root was defined by *Mycobacterium smegmatis.* Nodes with bootstrap confidences > 80 are indicated with gray circles, and precise bootstrap values are found in Figure S4A. The colored bars in the outer ring indicate genetic contributions from different ancestral populations as inferred by Structure analysis. The shading of the inner ring indicates the latitude from which each strain was isolated according to the scale provided. The isolation site for each strain can be determined by isolate names as indicated in Table 1.

**Table 1.**
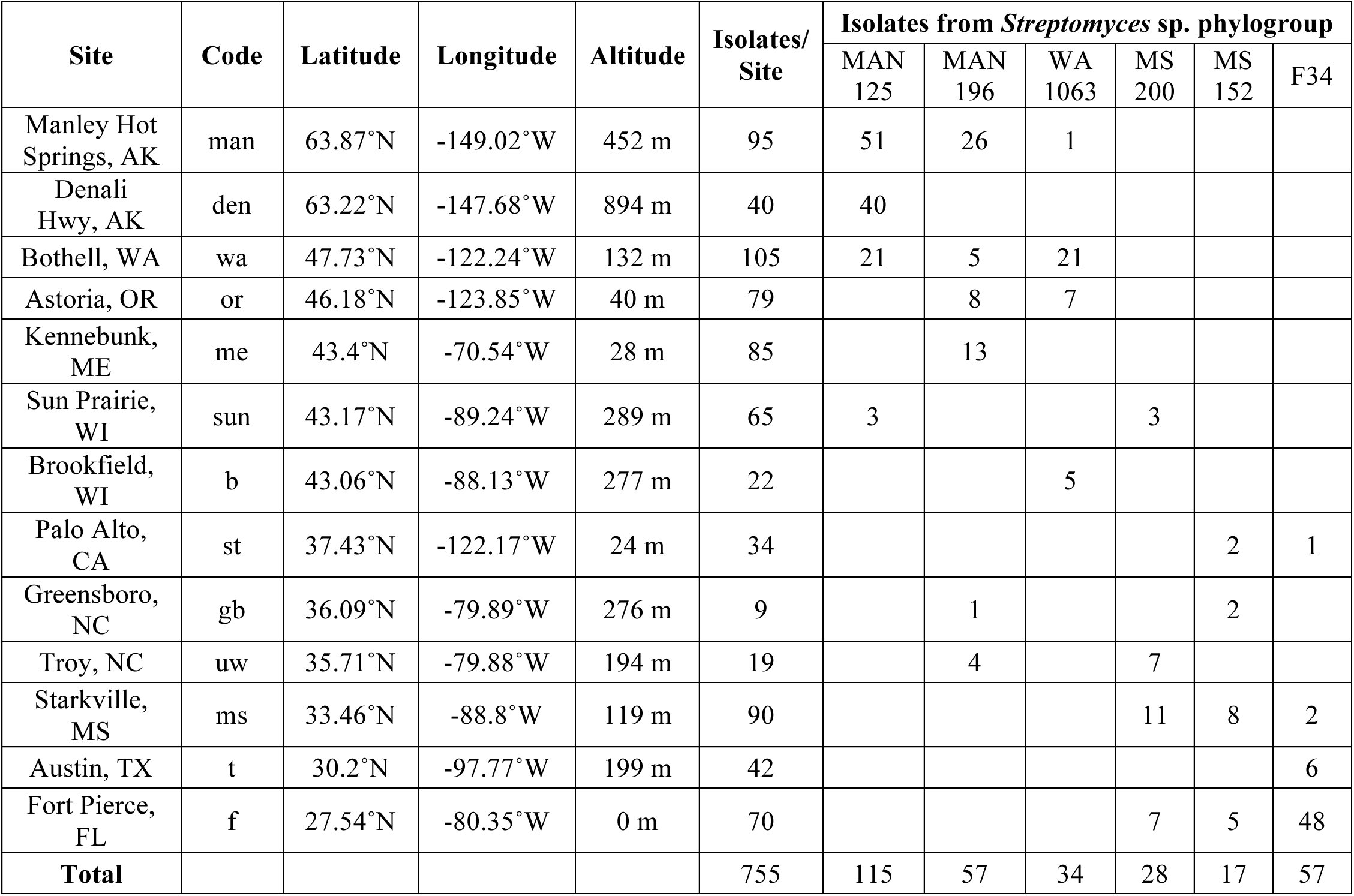
A total of 755 *Streptomyces* strains were isolated from 13 sites, and 308 of these strains were found to represent the six targeted phylogroups. The numbers of isolates per site that belonged to each of our target phylogroups is indicated. Isolate names begin with a letter code referring to the sample site.

**Table 2.** MLSA was performed on a total of 17-47 strains from each phylogroup. Summary statistics for concatenated MLSA nucleotide sequences were determined as described in methods. Phylogroup average latitude (Ave Lat) was determined using the total number of isolates per phylogroup provided in Table 1. The standard index of association is zero for a population in linkage equilibrium.

| Phylo-group | Ave Lat | n | Haplo-types | Min ANI | S | $\pi$ | Tajima's D | $\theta_w$ | $\rho$ | $\rho/\theta_w$ | $I_A$ | $\Phi_w$ |
| --- | --- | --- | --- | --- | --- | --- | --- | --- | --- | --- | --- | --- |
| MAN125 | 60.15°N | 45 | 25 | 99.5 | 23 | 0.0026 | 0.53 | 0.0023 | 0.0094 | 4.18 | 0.09 | 2.95E-01 |
| MAN196 | 52.84°N | 47 | 26 | 98.5 | 65 | 0.0075 | 0.67 | 0.0063 | 0.00043 | 0.068 | 0.49 | 4.24E-02* |
| WA1063 | 47.20°N | 19 | 13 | 98.1 | 63 | 0.0052 | -1.35 | 0.0077 | 0 | 0 | 0.43 | 4.33E-03 |
| MS200 | 33.58°N | 27 | 21 | 98.4 | 93 | 0.0088 | -0.58 | 0.0103 | 0.0017 | 0.17 | 0.39 | 1.33E-03** |
| MS152 | 32.50°N | 17 | 15 | 97.6 | 96 | 0.0107 | -0.51 | 0.012 | 0.0013 | 0.11 | 0.35 | 1.078E-01** |
| F34 | 28.20°N | 36 | 22 | 97.7 | 93 | 0.0109 | 0.52 | 0.0096 | 0.0011 | 0.11 | 0.46 | 1.79E-01** |

Ave Lat: average latitude of phylogroup members
n: number of isolates subsampled for MLSA
Min ANI: minimum percent pairwise average nucleotide identity (ANI) within phylogroup members
S: segregating sites
$\pi$ : per site nucleotide diversity
$\theta_w$ : per site Watterson's theta ( $2N_e\mu$ )
$\rho$ : per site rate of recombination/gene conversion ( $2N_e r/2$ )
$\rho/\theta_w$ : relative rate of recombination to mutation
$I_A$ : standard index of association

Four of the phylogroups do not match any *Streptomyces* species described in PubMLST (less than 97% ANI across MLSA loci, Figure S4A) (Jolley *et al*., 2004). While the other two, phylogroups WA1063 and MS152 belong to the *S. griseus* species cluster (Rong and Huang, 2010). Isolates in phylogroup WA1063 share greater than 99% MLSA identity with *S. anulatus* and *S. praecox* and form a monophyletic clade with these type strains (Figure S4A). Isolates in phylogroup MS152 share greater than 98% MLSA identity with *S. mediolani, S. albovinaceus*, and *S. griseinus* and form a paraphyletic clade that includes these type strains (Figure S4A).

We find evidence of horizontal gene transfer consistent with previous observations of *Streptomyces* species (Doroghazi and Buckley, 2010; Andam *et al*., 2016a). There is significant phylogenetic incongruence between MLSA loci (Figure S4, Table S2), suggesting that inter-species horizontal gene transfer has shaped the phylogeny of these groups. The six phylogroups exhibit evidence of population structure with each phylogroup composed of 3.2 ± 0.8 (mean ± s.d.) subpopulations and with evidence for admixture (Figure 1). Evidence of admixture suggests horizontal gene transfer within phylogroups and is consistent with previous evidence of gene exchange in *Streptomyces* (Doroghazi and Buckley, 2010). Evidence of recombination within phylogroups MAN196, MS200, MS152, and F34 (PHI test, *p* < 0.05, Table 2) further supports the conclusion of gene exchange within populations. Furthermore, the standard index of association (I_A_) is 0.09 for MAN125 suggesting a freely recombining population in linkage equilibrium (Table 2). In addition, phylogroups WA1063 and MS152 share two identical *atpD* alleles (Figure S4B), but it is not clear whether these alleles are shared as a result of contemporary horizontal gene transfer or vertical inheritance from the most recent common ancestor of the two clades. The latter explanation is more parsimonious given the low level of polymorphism between phylogroups

WA1063 and MS152. We do not observe evidence of inter-group horizontal gene transfer between these six phylogroups.

### Evidence for dispersal limitation

Strains of the six phylogroups were obtained from soil samples from 13 sites of diverse geographic origin (Table 1). Each phylogroup was detected in 4.2 ± 0.4 sites, and this distribution differs significantly from expectations for a random distribution of strains across sites (permutation test, *p* < 0.0005), thereby rejecting the hypothesis of panmixia (i.e. the ability of organisms to move freely across habitats). Each phylogroup subpopulation was observed in 2.2 ± 0.9 sites (mean ± s.d.), and this value is lower than expected if subpopulations are randomly distributed across the sites occupied by each phylogroup (permutation test, *p* < 0.001). These results indicate that phylogroup distribution is constrained geographically and that phylogroups have subpopulation structure that is also geographically explicit.

The geographic distribution of *Streptomyces* alleles indicates dispersal limitation. Identical alleles are shared among phylogroup members across each phylogroup’s geographic range, which can exceed 5,000 km (Figure 2, Table 1). However, dissimilarity in allele composition increases with geographic distance, and this result is significant (Bray-Curtis dissimilarity, Mantel r = 0.29, *p* = 0.005) (Figure S5). Hence, alleles are more likely shared between geographically similar sites indicating dispersal limitation with potential for long range dispersal. This result is significant for all individual loci except *recA* (Bray-Curtis dissimilarity, *aptD* Mantel r = 0.31, *p* = 0.004; *gyrB* r = 0.22, *p* = 0.031; *recA* r = 0.16, *p* = 0.088; *rpoB* r = 0.27, *p* = 0.004; *trpB* r = 0.19, p = 0.047). In addition, all MLSA haplotypes (Figure 1, Figure 3) are unique to a single site, with the sole exception being a haplotype from phylogroup MS200 which is observed in both MS and WI (Sun Prairie).

**Figure 2.**
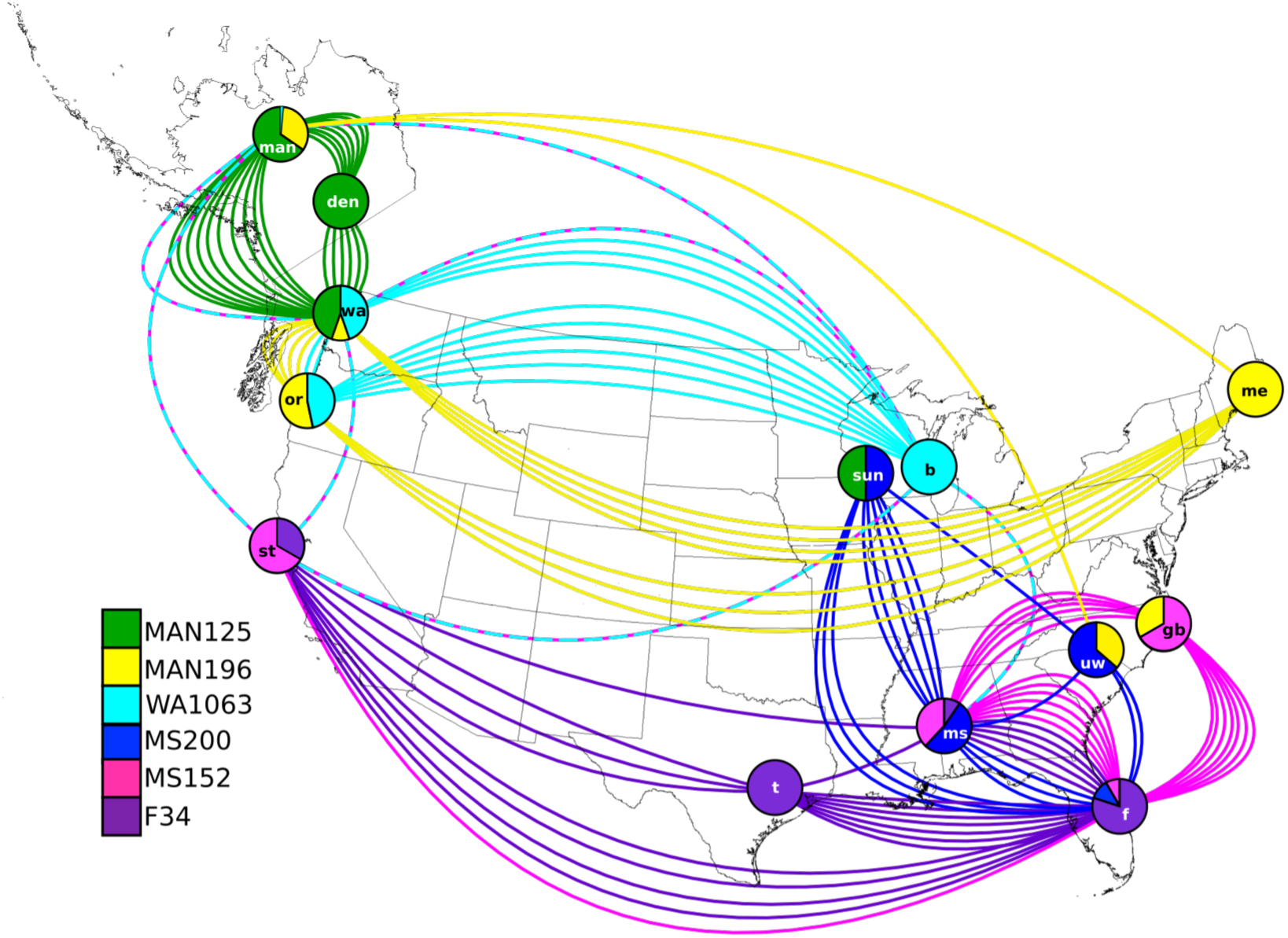
Circles depict sample sites and are labeled according to site code (Table 1). The relative abundance of each phylogroup at each site is indicated by color according to the legend, with raw counts provided in Table 1. Solid colored lines represent identical alleles shared by phylogroup members subsampled for MLSA across sites. Dashed multicolored lines depict identical *atpD* alleles shared by strains in phylogroups WA1063 and MS152 (Figure S4B).

Analysis of haplotype distribution is consistent with diversification due to dispersal limitation. We used nested clade analysis (NCA) combined with a rigorous statistical framework to evaluate population structure and demography (see Experimental Procedures). NCA establishes significant phylogeographic inferences for phylogroups MAN196, MAN125, WA1063, and MS152 (Figure 3) but not for MS200 and F34. Nested clade phylogeographic inference postulates potential evolutionary and historical demographic processes that support extant patterns of diversity and biogeography (see legend of Figure 3). For instance, population subdivision of MAN125 across the Pacific Northwest (Figure 3A) and MAN196 between Maine and the Pacific Northwest (Figure 3B) is consistent with restricted gene flow due to historical long distance dispersal events. Likewise, population subdivision of MS152 and between the Southeast (MS and FL) and CA (Figure 3C) and WA1063 between WI (Brookfield) and OR (Figure 3D) is consistent with allopatric fragmentation.

**Figure 3.**
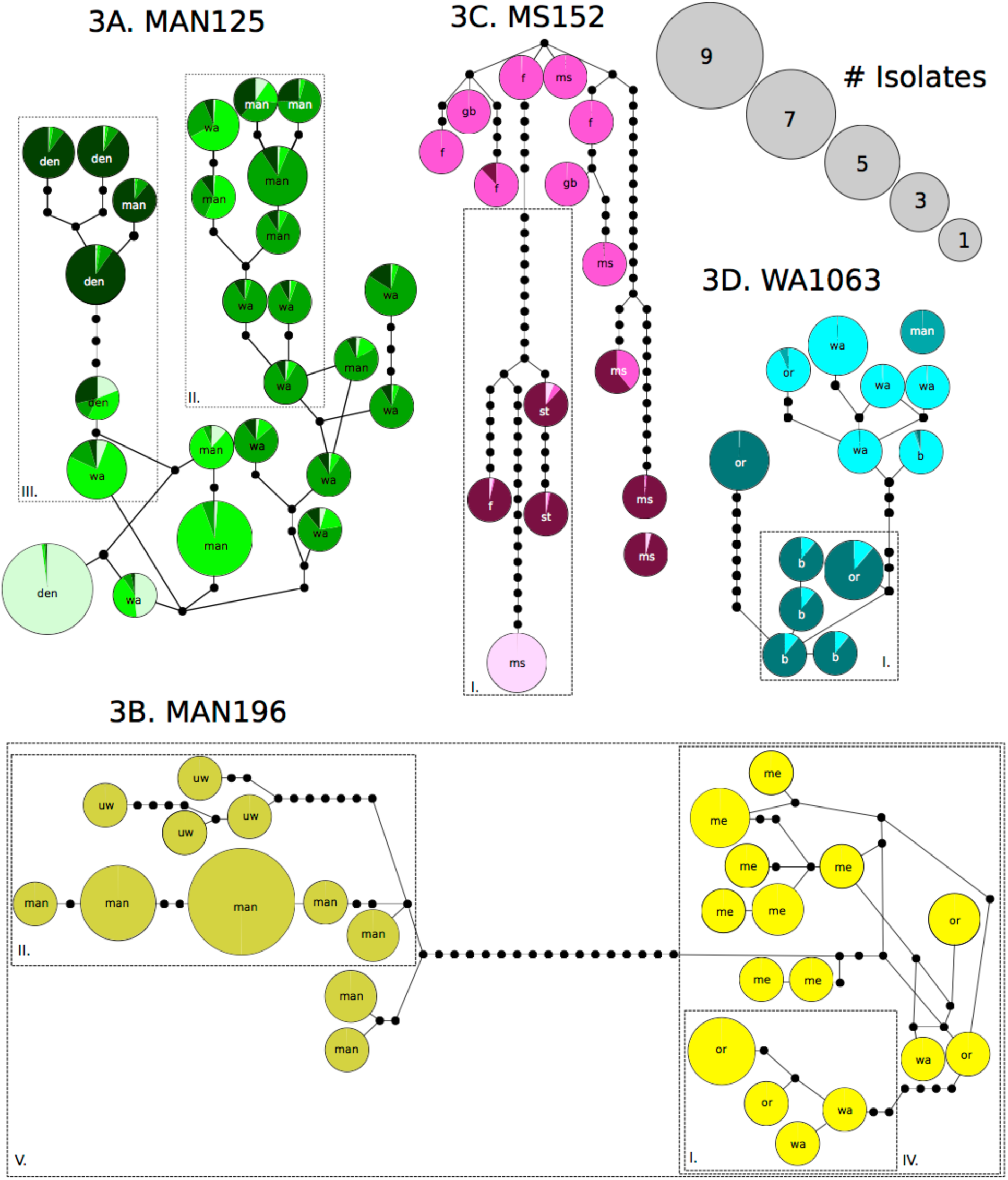
Haplotype networks for phylogroups MAN125 (3A), MAN196 (3B), MS152 (3C), WA1063 (3D). Circles represent MLSA haplotypes whose radius is proportional to the number of isolates having that haplotype, and colors correspond to strain ancestry and subpopulation affiliation as determined by Structure analysis and defined in Figure 1. Haplotypes are labeled with a letter code referring to sample site location as indicated in Table 1 (all haplotypes for MAN125, MAN196, MS152, and WA1063 are found exclusively at a single site). Black circles represent unsampled, inferred haplotypes with each circle designating a single nucleotide polymorphism. The length of edges between nodes is uninformative. Dashed rectangles encompass clades that have significant phylogeographic inferences from nested clade analysis, as described in methods. Roman numerals correspond to the following inferences: I. Allopatric fragmentation; II. Long distance colonization and/or past fragmentation; III. Restricted gene flow with isolation by distance; IV. Restricted gene flow but with some long-distance gene flow over intermediate ranges not occupied by the species; or past gene flow followed by extinction of intermediate populations; V. Contiguous range expansion.

### Latitudinal diversity gradient

The distribution and diversity of the phylogroups reveals a latitudinal diversity gradient. Strains from MAN125, MAN196, WA1063 occur mostly north of 40°N latitude, while strains from MS200, MS152, F34 occur mostly south of this latitude (Table 1, Figure 2). This pattern of North/South partitioning is significant for each phylogroup when evaluated against the expectation of a random distribution across sites (permutation test, *p* < 0.01 for each phylogroup after Bonferroni correction). Furthermore, partial Mantel tests were performed to evaluate the latitudinal and longitudinal vector components of geographic distance in relation to the allele composition of sites. There remains a significant relationship between allele composition and geographic distance when we control for longitude, (Mantel r = 0.23, *p* = 0.022), but this relationship is no longer significant when we control for latitude (Mantel r = 0.15, *p* = 0.12). This result indicates that allele composition changes more across latitude than it does across longitude. The latitudinal partitioning of alleles can be readily observed in the pattern of allele sharing between sites (Figure 2). Finally, we also observed a significant relationship between per site nucleotide diversity of phylogroup MLSA loci and the average latitude of sites in which they are found (R = −0.91, *p* = 0.012; Figure 4). This result indicates phylogroups recovered from lower latitudes have higher genetic diversity than those recovered from higher latitudes.

**Figure 4.**
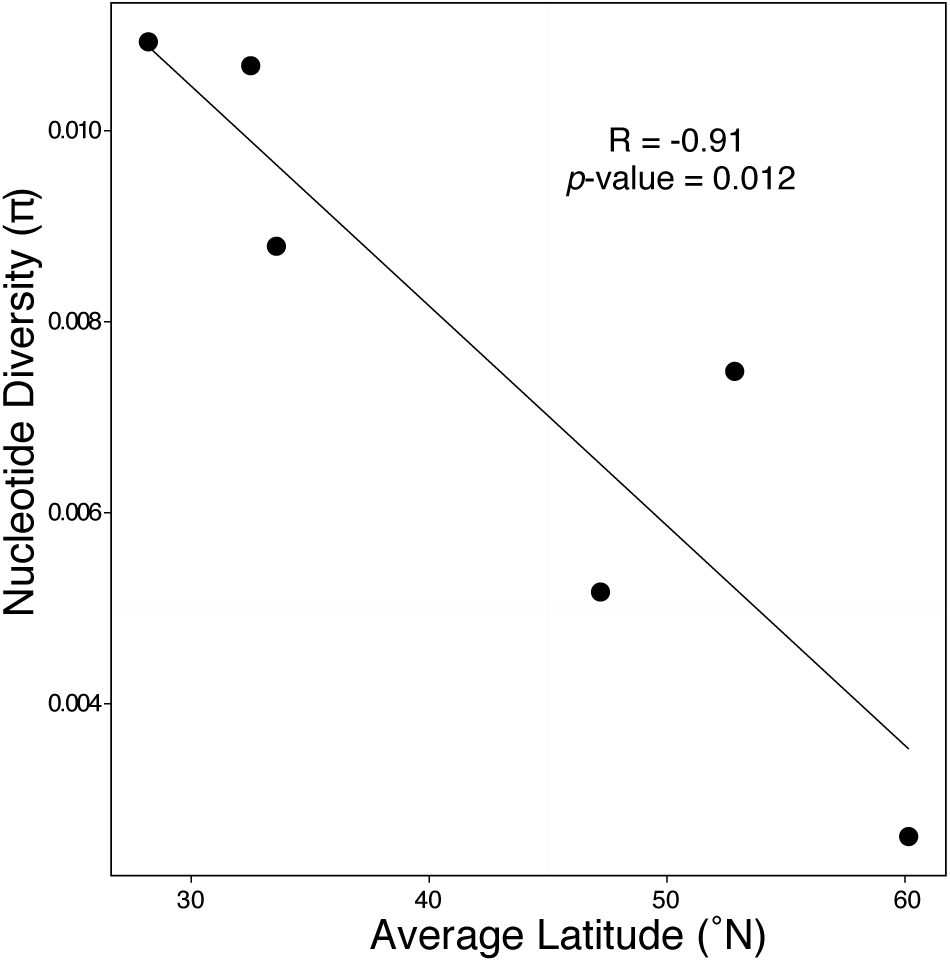
Average latitude was determined using a weighted average based on the total number of isolates per site as indicated in Table 1. Nucleotide diversity was calculated using concatenated MLSA loci and is expressed per site.

## Discussion

We used population genetic approaches to analyze spatial patterns of genetic diversity for six *Streptomyces* phylogroups isolated from geographically disparate but ecologically similar sites across the United States (Table 1). The distribution of phylogroups is nonrandom and likely dispersal limited (Figure S5) with phylogroups inhabiting geographic ranges defined by latitude (Table 1, Figure 2). In addition, the genetic diversity of phylogroups is negatively correlated to the latitude from which they were isolated (Figure 4). These findings suggest that there are latitudinal barriers to dispersal, and that patterns of *Streptomyces* biogeography result from dispersal limitation and regional diversification due to genetic drift. Furthermore, these results are consistent with the hypothesis that historical demographic processes have influenced the contemporary biogeography of *Streptomyces.*

The phylogroups we describe are coherent phylogenetic groups that approximate biological populations (Figure S2). The members of each phylogroup share a distinguishable geographic range (Table 1, Figure 2), a recent common ancestor (Figure 1), and greater than 97% ANI across MLSA loci (Table 2). Despite geographic and genetic subpopulation structure (Figure 1), phylogroup members frequently share identical alleles across demes. It is worth noting that since all of the strains examined share > 97% SSU rRNA gene identity, these geographic patterns would not be detected using standard SSU rRNA analyses methods (Figure S1). We infer the presence of recombination within all phylogroups using both nucleotide polymorphism and phylogenetic methods (Table 2, Table S2).

Regional patterns of biogeography among phylogroups are consistent with limitations to dispersal and gene flow. Allopatric processes like genetic drift can drive diversification between populations that are geographically isolated. The geographic distribution of our phylogroups is nonrandom, and we find regional subpopulation structure within phylogroups (Figure 1, Figure 3). Although we find identical alleles in sites thousands of kilometers apart (Figure 2, Figure S4), MLSA haplotypes are not shared across sites (with the single exception of a haplotype shared between MS and WI) (Figure 3). We also observe a significant distance decay relationship for MLSA allele composition and geographic distance between sites (Figure S5). Distance decay relationships driven by neutral processes can be challenging to identify given that environmental variables are often spatially structured (Nekola and White, 1999). However, Andam *et al.* (2016b) shows that *Streptomyces* community phylogenetic diversity across sites correlates significantly with latitude and temperature but not with soil pH, soil organic matter, or rainfall. Hence, it is unlikely that these coarse environmental variables determine regional population structure. This data implies that while gene flow is moderate across the geographic range of a phylogroup, dispersal limitation and genetic drift create appreciable regional population structure.

*Streptomyces* phylogroup diversity is consistent with a latitudinal diversity gradient. While the classical description of a latitudinal diversity gradient defines diversity at the level of species richness, these diversity gradients are also apparent at the level of intra-species genetic diversity (Hadly, 2013). We find that latitude is a significant predictor of gene flow (Figure 2). Furthermore, intra-phylogroup nucleotide diversity has a significant negative relationship with average latitude (Figure 4), which is congruous with the latitudinal diversity gradient observed for diverse macroorganisms (Hadly, 2013; Hillebrand, 2004). There is conflicting evidence for latitudinal diversity gradients among microorganisms. Evidence for microbial latitudinal diversity gradients comes from marine systems (Fuhrman *et al*., 2008; Sul *et al*., 2013; Swan *et al*., 2013), with contrary evidence obtained in terrestrial systems (Neufeld and Mohn, 2005; Chu *et al*., 2010). However, most analyses of terrestrial bacterial biogeography are derived from analyses of SSU rRNA genes, and we show that analyses of SSU rRNA genes lack the sensitivity needed to detect the biogeographic patterns that we observe for *Streptomyces* (Figure S1).

Several hypotheses have been advanced to explain the formation of latitudinal diversity gradients (Wiens and Donoghue, 2004; Mittelbach *et al*., 2007). Ecological hypotheses posit that factors such as carrying capacity, productivity, and niche availability vary across latitude and that these factors impose constraints on biodiversity (Currie *et al*., 2004; Mouchet *et al*., 2015). Evolutionary hypotheses invoke the positive relationship between temperature and the kinetics of metabolism to predict that evolutionary rates and cladogenesis correspond with temperature (Allen *et al*., 2002). Historical hypotheses propose that the latitudinal diversity gradient is the product of historical geological, ecological, or demographic events that have influenced dispersal and diversification (Wiens and Donoghue, 2004; Stevens, 2006). For example, the influence of Pleistocene glacial events on the biogeography of diverse species of terrestrial and aquatic plants and animals is well documented, with postglacial range expansion giving rise to the latitudinal partitioning of populations and species, and resulting in decreased molecular variation of northern lineages relative to southern ancestral lineages (Soltis *et al*., 1997; Bernatch and Wilson, 1998; Conroy and Cook, 2000; Milá *et al*., 2006; Maggs *et al*., 2008; Wilson and Veraguth, 2010). There is also evidence that Pleistocene glaciation events have impacted both microbial communities (Eisenlord *et al*., 2012) and populations (Kenefic *et al*., 2009; Mikheyev *et al*., 2008).

The biogeography of our *Streptomyces* phylogroups is consistent with the hypothesis that historical demography and dispersal limitation have produced the latitudinal diversity gradient that we observe. For instance, MAN125 is nearly in linkage equilibrium (I_A_ = 0.09) though its members span a geographic range of over 2,000 km across the Pacific Northwest. Similar patterns of recombination have also been observed in a *S. pratensis* population which spanned 1,000 km across sites present in North Carolina and northern New York (Doroghazi and Buckley, 2010 and 2014). The observation of linkage equilibrium indicates that either the population lacks contemporary barriers to gene flow or the population has experienced a recent historical demographic expansion. Coupled with evidence of limitations to gene flow at regional scales (Figure 2, Figure S5), the most parsimonious explanation for these conflicting observations is a recent historical demographic range expansion. Conversely, latitudinal gradients in marine bacterial diversity have been attributed to correlations between temperature and the kinetics of metabolism (Fuhrman *et al*., 2008). This kinetic effect has been hypothesized to increase evolutionary tempo and speciation rates in tropical latitudes and would be expected to generate latitudinal gradients of both species diversity and nucleotide diversity (Allen *et al*., 2002). Since latitude is correlated with temperature, we cannot completely dismiss the influence of kinetics as a cause of the intra-group latitudinal gradient of genetic diversity in terrestrial *Streptomyces*. However, the kinetic effects of temperature cannot account for partitioning of gene flow across latitude, while in contrast, this is a specific prediction of the historical demography hypothesis. It is possible, however, that unappreciated ecological variables, such as the species composition of perennial grass communities, could shape the diversity gradient. Ultimately, it is likely that latitudinal diversity gradients can arise from a combination of ecological, evolutionary, and historical processes that vary in their relative influence with respect to different species and different habitats.

Evidence for a latitudinal diversity gradient coupled with evidence of contemporary dispersal limitation following historical demographic expansion, while not conclusive, suggests that phylogenetic niche conservatism has contributed to the formation of the *Streptomyces* latitudinal diversity gradient. Phylogenetic niche conservatism can cause diversity gradients when contemporary species distributions are determined by historical climate regimes (Wiens and Donoghue, 2004; Stevens, 2006). Climactic regimes oscillate widely across geologic time scales and these historical changes in climate produce demographic phenomena that impact the evolutionary dynamics of species. In particular, the genetic consequences of glaciation events are described in depth by Hewitt (1996, 2000, and 2004). The population structure we observe in our *Streptomyces* phylogroups is consistent with the effects of post-glacial demographic range expansion followed by dispersal limitation and regional diversification. Dispersal limitation may occur following range expansion as a result of density dependent blocking or by adaptive barriers that arise after the colonization of new habitat. One of the expectations of post-glacial expansion is “southern richness versus northern purity” (Hewitt, 2004). This is evident in the negative correlation we observe between latitude and the nucleotide diversity of phylogroups (Figure 4). Similar relationships between intraspecific nucleotide diversity and average latitude as result of post-glacial colonization are evident in other systems (Bernatchez and Wilson, 1998). Williams *et al.* (1998) justifies 40°N latitude as approximating late Pleistocene glacial and non-glacial regions with respect to species distributions in North America. Hence, the latitudinal delineation of allele distributions for *Streptomyces* phylogroups roughly corresponds to the extent of ice coverage during the late Pleistocene (Figure 2), which suggests historical population expansion from lower to higher latitudes.

Haplotype distributions of phylogroups MAN125, MAN196, WA1063, and MS152 are consistent with allopatric diversification resulting from dispersal limitation (Figure 3). Haplotype nested clade analysis (NCA) predicts that historical dispersal events and range expansion across northern regions resulted in limits on dispersal during intermediate timescales, allowing genetic drift to create the phylogroup population structures observed today. There is moderate criticism of NCA (Knowles and Maddison, 2002; Nielsen and Beumont, 2009) due to the subjective nature of inferring historical processes and the wide potential for stochastic processes creating similar patterns of biogeography. Yet these tools can provide useful hypotheses. Northern phylogroups MAN125 and MAN196 share a common ancestor with southern phylogroups MS200 and F34 (Figure 1). The contemporary population structure of MAN125 and MAN196 is consistent with a historical range expansion from a common ancestor shared by both clades (Figure 1, Figure 3). Further population structure within each phylogroup likely resulted from barriers to gene flow and historical dispersal events across the Pacific Northwest for MAN125 (Figure 3A) and between Maine and the Pacific Northwest for MAN196 (Figure 3B). Analysis of haplotype distribution within WA1063 and MS152 is also consistent with diversification of populations as a result of gene flow limitation between the Midwest and Pacific Northwest for WA1063 (Figure 3D) and between the Southeast and West for MS152 (Figure 3C).

Phylogroups WA1063 and MS152 share a recent common ancestor (Figure 1, Figure S2) and also share two identical alleles at the *atpD* locus (Figure S4B). These phylogroups have distinct, nonoverlapping geographic ranges with WA1063 found in higher latitudes and MS152 in lower latitudes (Table 1, Figure 2). WA1063 and MS152 have 0.0205 net nucleotide substitutions per site across concatenated MLSA loci. We evaluate the possible time range for divergence between WA1063 and MS152 by extrapolating very roughly from the nucleotide substitution rate (μ = 4.5×10^−9^) and generation time (100-300 generations per year) for *E. coli* (Ochman *et al*., 1999), since corresponding values are not available for *Streptomyces* or their relatives. Based upon these gross approximations, we would estimate that WA1063 and MS152 diverged 15,000-50,000 years ago, corresponding to events in the late Pleistocene (Clayton *et al*., 2006). Hence, it is likely that the identical *atpD* alleles found in both WA1063 and MS152 were inherited from a shared ancestral population (Figure S3B) as opposed to inheritance from contemporary gene exchange.

Through population genetic analysis of six *Streptomyces* phylogroups, we find evidence for dispersal limitation associated with geographically explicit patterns of gene flow which manifest in a latitudinal gradient of nucleotide diversity. Furthermore, these data support the hypothesis that historical demographic processes influence the contemporary biogeography of *Streptomyces.* Due to their spore forming capabilities and potential for long range dispersal, *Streptomyces* are an ideal system for assessing limits on gene flow among terrestrial bacteria. Future research should seek to determine the degree to which dispersal limitation is due to limits on spore mobility, density dependent blocking, or a result of adaptive constraints relating to phylogenetic niche conservatism. A better understanding of *Streptomyces* biogeography and the evolutionary forces that govern *Streptomyces* diversification may ultimately assist in the discovery of novel genetic diversity and possibly novel antibiotics within this genus.

## Experimental Procedures

### Strain isolation and DNA extraction

We previously assembled a culture collection of more than 1,000 *Streptomyces* from 15 sites across the United States with soil sampled at 0-5 cm depth (Andam *et al*., 2016b). Sites were selected to represent a narrow range of ecological characteristics including meadow, pasture, or native grasslands dominated by perennial grasses and having moderately acidic soil (pH: 6.0 ± 1.0; ave. ± s.d.). Strains were isolated using uniform conditions and this will select for strains having similar physiological characteristics. The analysis of physiologically similar strains from ecologically similar sites improves our ability to detect biogeographical patterns that result from drift by minimizing the importance of selection (as reviewed by Hanson *et al*., 2012). Soil was air dried, and *Streptomyces* strains were isolated on glycerol-arginine agar plates of pH 8.7 containing cycloheximide and Rose Bengal (El-Nakeeb and Lechevalier, 1963; Ottow, 1972) as previously described (Doroghazi and Buckley, 2010). Genomic DNA was extracted for each isolate from purified cultures, which were grown by shaking at 30°C in liquid yeast extract-malt extract medium (YEME) containing 0.5% glycine (Kieser *et al*., 2000), by using a standard phenol/chloroform/isoamyl alcohol protocol (Roberts and Crawford, 2000).

The gene encoding the RNA polymerase beta-subunit *(rpoB)* provides a robust, advantageous alternative to the SSU rRNA locus for phylogenetic analyses of the genera *Streptomyces* (Kim *et al*., 2004). We previously assessed genetic diversity of our culture collection using partial *rpoB* sequences clustered at 0.01 patristic distances with RAMI (Pommier *et al*., 2009), and using this approach we identified 107 species-like phylogenetic clusters, or phylogroups (Andam *et al*., 2016b). We selected six of these phylogroups for further analysis. These six phylogroups had the highest numerical abundance and widest geographical distribution in our isolate collection, representing 308 strains isolated across 13 sites (and representing 308 of the 755 strains isolated from these 13 sites). MLSA was performed on 17-47 isolates from each phylogroup. Phylogroup names are capitalized and based on a representative isolate; for example, phylogroup WA1063 is named for *Streptomyces* sp. wa1063. Isolates are identified with a lowercase letter code indicating site of origin followed by a strain number; for example, isolate wa1063 is strain 1063 isolated from Washington (WA) state.

### Multilocus sequence analysis (MLSA)

We adapted the MLSA scheme developed for *Streptomyces* by Guo *et al.* (2008), which targets the five housekeeping loci *atpD, gyrB, recA, rpoB*, and *trpB* as described in Doroghazi and Buckley (2010) (Table S3). The V2 and V2 regions of SSU rRNA sequences were amplified using universal primers 8F and 1492R (Table S3). Reactions for Sanger sequencing were performed using forward primers for all loci except *rpoB*, for which the reverse primer was used. Trace files were uniformly screened using CAP3 (Huang and Madan, 1999), and sequences were trimmed as to discard nucleobases with a Phred quality score below 23. Sequences were aligned using MUSCLE (Edgar, 2004), trimmed to 431 bp, 415 bp, 446 bp, 557 bp, and 489 bp, for genes *atpD*, *gyrB*, *recA*, *rpoB*, *trpB*, respectively, and concatenated consistently with the genomic order in *Streptomyces coelicolor* A3(2). SSU rRNA sequences were trimmed to 357 bp, creating an alignment spanning the V1 and V2 regions. Gene sequences are available on GenBank with accession numbers KX110408-KA111380.

Good’s coverage estimation and haplotype rarefaction was determined using mothur (Schloss *et al*., 2009). DNA polymorphism statistics including, number of segregating sites, nucleotide diversity, and Tajima’s D, were determined with DnaSP v5 (Librado and Rozas, 2009) and LDhat (McVean *et al*., 2002). Population scaled mutation rates (Watterson's theta; θ_w_= 2N_e_μ), recombination or gene conversion rates (ρ=2N_e_r/2), and relative rates of recombination (ρ=/θ=_w_) were estimated using LDhat (McVean *et al*., 2002) and are expressed per nucleotide site. The standard index of association (I_A_) was calculated from allelic data with LIAN v3.5 (Haubold and Hudson, 2000) using the Monte-Carlo test and 100 iterations. The pairwise homoplasy index (PHI) statistic was determined using PhiPack (Bruen *et al*., 2006), and statistical significance was evaluated under a null hypothesis of no recombination. Sequence identity across phylogroups was calculated with mothur (Shloss *et al*., 2009)

### Phylogenetic reconstruction

Maximum likelihood (ML) trees were constructed from the nucleotide sequences of individual and concatenated MLSA loci using the generalized time reversible nucleotide substitution model (Tavaré, 1986) with gamma distributed rate heterogeneity among sites (GTRGAMMA) supported in RAxML v7.3.0 (Stamatakis, 2006). Bootstrap support was determined for the highest-scoring ML tree of 20 iterations, and the number of bootstrap replicates was determined using the extended majority rule (autoMRE) convergence criteria (Pattengale *et al*., 2010). Root placement is defined by *Mycobacterium smegmatis.* Significant phylogenetic incongruence between loci was determined using the Shimodaira-Hasegawa test (Shimodaira and Hasegawa, 1999) implemented in the R package phangorn (Schliep, 2011).

### Population structure

Concatenated MLSA sequences were analyzed using Structure v2.3.3 (Pritchard *et al*., 2000) to examine population affiliation, subdivision, and admixture within and between phylogroups. Structure was run using an admixture model with a burn-in length of 1.0E^6^ and 3.0E^6^ replicates. The most probable number of sub-populations (k) was evaluated with 10 independent runs and chosen using the Evanno method (Evanno *et al*., 2005), with k = 1 through k = 6 within phylogroups and through k = 8 between phylogroups, implemented by Structure Harvester (Earl and vonHoldt, 2012). After choosing the most probable k-value, the program Clumpp was used to permute outputs of the independent runs (Jakobsson and Rosenberg, 2007).

### Patterns of dispersal and gene flow

We used permutation tests to evaluate whether phylogroup distribution across sites could be explained by panmixia. We compared the observed distributions to those expected under a random distribution model using 1,000 permutations to assess significance. The null model for the permutation test assigned strains to sites as a random draw without replacement from the OTU table while holding the number of strains sampled at each site to be invariant. In addition, correlations between geographic distance and allele composition between sites were assessed using Mantel and partial Mantel tests (Mantel, 1967; Smouse *et al*., 1986). These tests were performed with the R package ecodist (Goslee and Urban, 2007) using the Pearson correlation method and 1,000 permutations. Bray-Curtis dissimilarity was calculated from allele composition across sites.

Haplotype networks were created using a statistical parsimony procedure (Templeton *et al*., 1987; Templeton *et al*., 1995) implemented in TCS v1.18 (Clement *et al*., 2000). Nested clade information was used to infer processes that could explain the geographic and genetic distribution of sequences using the program GeoDis v2.2 (Posada *et al*., 2000). Both TCS v1.18 and GeoDis v2.2 were performed in ANeCA (Panchal, 2007).

## Acknowledgements

This material is based upon work supported by the National Science Foundation under Grants No. DEB-1050475 and DEB-1456821. We would like to thank Ashley Campbell for her help in generating MLSA sequences.

